# Tracking *Helicoverpa zea* (Lepidoptera: Noctuidae) flight behavior with infrared camera around pheromone trap

**DOI:** 10.1101/2024.12.07.627224

**Authors:** Christophe Duplais, Chris Roh

## Abstract

Understanding the flight behavior of nocturnal insect pests is essential for designing effective trapping systems and improving integrated pest management (IPM). This study used infrared (IR) reflectance and IR camera to analyze the flight behavior of corn earworm (*Helicoverpa zea*) around Scentry Heliothis pheromone trap, focusing on approach, escape, and capture rates. Video analysis revealed low average catch rate of 25% for a total of 48 approaches from 7 p.m. to 4 a.m., as many moths approached the lure but escaped without being trapped. The study identified critical behavioral patterns, such as upwind approach, downwind horizontal escape, or vertical ascending leading to capture. These findings suggest that positioning lure closer to the trap entrance could significantly improve the effectiveness of Scentry Heliothis trap. Additionally, the study highlights the importance of developing a comprehensive database of nocturnal insect behavior around trapping systems. This knowledge can be used to refine trap design for specific insect pest and to update catch count thresholds to improve the effectiveness of insecticide spray programs in precision agriculture. This work demonstrates the potential of IR cameras as a simple, commercially available, and affordable tool for studying nocturnal insect pest activity and flight behavior, with the potential to enhance pest management strategies in agriculture.

## Introduction

Understanding the flight behavior of insect pests around trapping systems is critical for designing effective monitoring and control strategies in agricultural systems (Miller et al., 2015). Trapping systems, including delta, sticky, funnel, bucket, Scentry Heliothis, and Hartstack traps, whether or not they employ a lure imbedded with female insect sex pheromone, are integral to integrated pest management (IPM) (Muirhead-Thomson, 1991; El-Sayed et al., 2006; Shelton and Badenes-Perez, 2006). Trapping systems providing valuable data on pest population dynamics, seasonal activity, and migration patterns, especially in the context of climate change (Lawton et al., 2022). However, the success of these systems depends on how well they align with the flight behavior and sensory physiology of the target insect species. Despite their importance, key aspects such as approach trajectories, trapping success rates, and avoidance maneuvers are poorly understood, limiting the optimization of trap selection and trap design. While pheromone-based traps are a cornerstone of IPM, their performance varies significantly due to incomplete understanding of nocturnal insect flight behaviors around trapping systems. This knowledge gap is particularly striking given the economic impact of moth pests, whose larvae cause billions of dollars in agricultural damage worldwide (Renault et al., 2022).

Although the increasing use of camera traps enables automatic detection and identification of captured insects (Cardim Ferreira Lima et al., 2020; Preti et al., 2021; Suto, 2022), traditional methods of studying insect flight around traps by direct observation are often limited by the absence of light at night during nocturnal insect activity. The inability to directly observe moth flight behavior hinders not only our ability to predict and explain why certain traps perform better for a given species but also our capacity to improve trap design to maximize insect capture. Infrared (IR) light, which emit IR light reflected off objects (IR reflectance) coupled with IR camera, differ from IR thermal camera, which detects heat emitted by animals and are significantly less sensitive. IR reflectance and cameras offer an affordable and practicable solution to these challenges. IR reflectance and cameras have been used in the labs (Fandino et al., 2019; Adam et al., 2021; Baleba et al., 2023), but field study insects with IR camera is limited to remote sensing with few practical applications (Rhodes et al., 2022). By capturing flight behavior during night time under moth-insensitive IR illumination, IR camera enables entomologist researchers and extension specialists to observe natural flight behavior around trapping systems without interfering with insect activity. This approach is particularly advantageous for nocturnal moths, whose behaviors are predominantly influenced by specific olfactory cues (female sex pheromone) in trapping systems at night.

This study leverages IR reflectance and cameras to address these gaps by analyzing the flight behavior of insect pests around trapping systems. By documenting approach trajectories, escape flight, and trapping outcomes, we aim to uncover key behavioral patterns that influence trap effectiveness. The insights gained will inform the development of more efficient trap designs tailored to the specific behavior of target pests, enhancing pest monitoring and control strategies across agricultural systems. Our initial findings focus on the flight behavior of the corn earworm (*Helicoverpa zea*) around the Scentry Heliothis trap. Corn earworm, a major insect pest in North America, lays eggs on corn silk and the larvae cause extensive damage to corn ears (Bohnenblust et al., 2013; Bibb et al., 2018; Reay-Jones, 2019; Britt et al., 2021). The sex pheromone mixture of (*Z*)-11-hexadecenal (Z11-Ald16) and (*Z*)-9-hexadecenal (Z9-Ald16) has proved to be highly effective as a lure (Sekul et al., 1975; Klun et al., 1980; Vetter and Baker, 1984), with the Hartstack trap demonstrating superior performance among all trap types tested (Gauthier et al., 1991; Latheef et al., 1993; Lopez et al., 1994; Guerrero et al., 2014). The significant difference in performance between the Heliothis Scentry trap and the Hartstack trap with higher number caught in the latter one, despite their similar conical design, is puzzling and suggests more investigation into the underlying factors driving this difference (Guerrero et al., 2014). Therefore, the results herein highlight potential flaws and offer actionable recommendations for improving trapping performance of Scentry Heliothis trap, laying the groundwork for enhanced IPM solutions in agriculture.

## Materials and Methods

### Setup with IR Camera and IR Light

The infrared camera, Phasm Cam, was purchased from GhostStop. The video resolution was set to 1080p HD (1920×1080 at 60 fps). Additional accessories included an infrared light (Orbo LN-5 or Andoer Mini IR Light) emitting at 850 nm, an external portable charger (50000 mAh), a 512GB Lexar E-Series Micro SD card, and a NEEWER T91 mini tripod, all purchased on Amazon. A hole has been drilled in the protective case to connect the Phasm Cam to the external battery overnight via the power cable. The IR light is also connected to the same external battery overnight. The connections are protected by electrical bandage in case of rain. The entire setup was acquired for under $300 (Figure 1A).

**Fig. 1.**
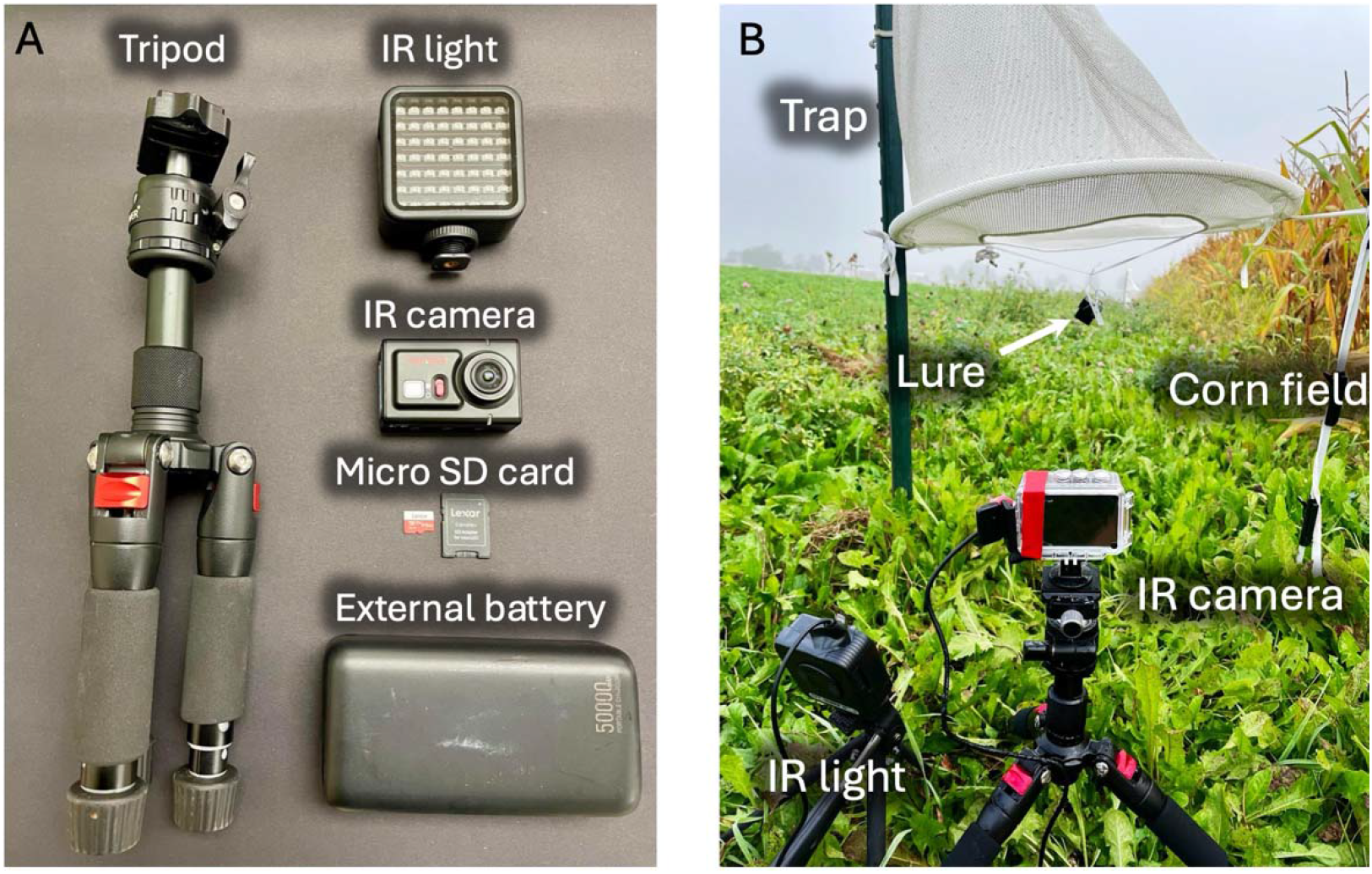
Equipment for overnight filming with an IR camera (A) and setup configuration in front of the trap and pheromone lure (B).

### Monitoring *H. zea* Flight Behavior and Activity

Two Scentry Heliopsis traps were set on the west side of a cornfield at Kime Farms near Geneva, NY (42.844275, -76.910826) on 19 September 2024. Each trap was equipped with either Hercon or Trece lure, both containing a blend of 97% (*Z*)-11-hexadecenal (Z11-16Ald) and 3% (*Z*)-9-hexadecenal (Z9-16Ald). These lure are comparable in terms of efficacy of trapping corn earworm (unpublished results).

### Data Analyzing

The .mov videos were analyzed using QuickTime Player software, with high-speed playback used to rapidly identify corn earworm approach events. This process allowed for the quantification of approaches that resulted in trapping versus non-trapping outcomes and the determination of the time of the approach. Photoshop was used to create the stacked video frames and ImageJ to create the colored flight trajectories. All successful and failed trapping videos can be found in Dryad data repository at http://datadryad.org/stash/share/E78OXHMsYtNc5rTXJ5RrazKB9zQSe8vJPae8rCdCBe8. Two short videos are also available in supporting files S1 and S2.

## Results

The equipment was placed in the field in front of each trap, as shown in Figure 1B. Five and seven corn earworms were captured in traps with Trece and Hercon lures, respectively. Video analysis revealed 28 approaches to the Trece lure and 20 approaches to the Hercon lure, resulting in capture rate of 18% and 35% per approach, respectively, and 25% on average. Combined data from both lures, presented in Figure 2, illustrating the distribution of observed behaviors (approach, capture, and escape) and reporting that corn earworm nocturnal activity spans from 7 p.m. to 4 a.m..

**Fig. 2.**
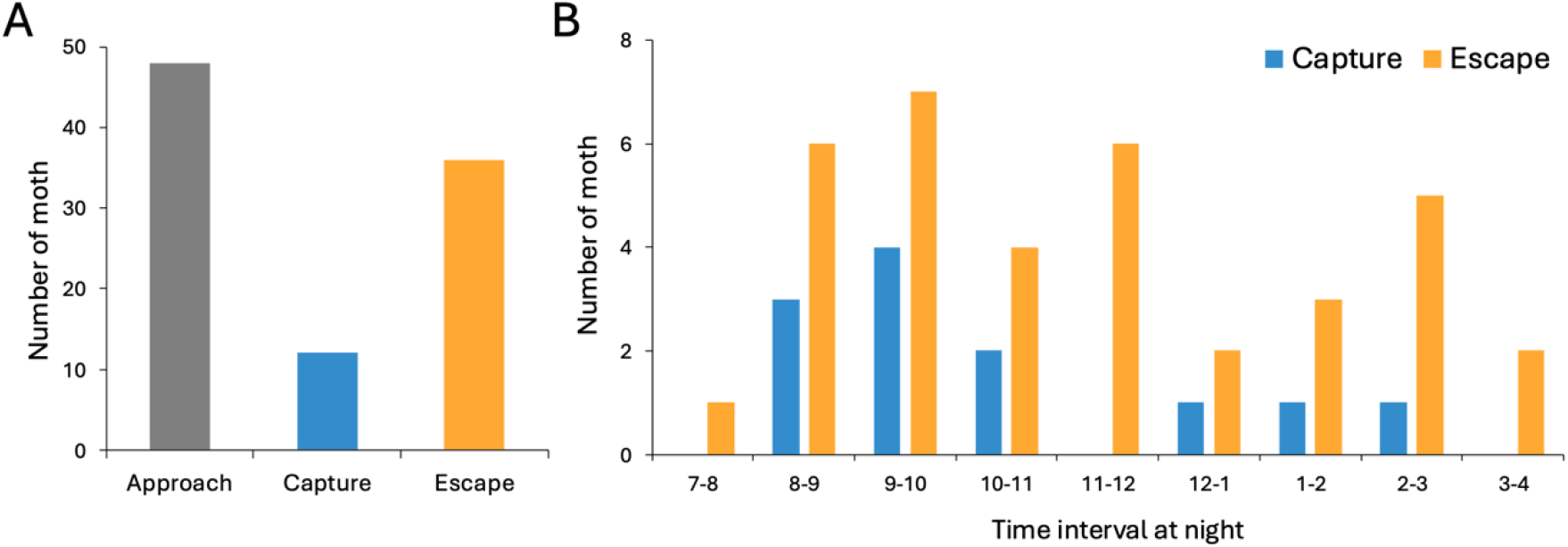
Behavioral analysis showing the total number of corn earworm approach, escape, and capture (A) during one-hour interval from 7 p.m. to 4 a.m. (B).

Video analysis shows that corn earworm approaches the lure closely for a few seconds, typically flying out upwind of the cornfield, guided by the pheromone plume emitted by the lure. Although corn earworm often positions just below the trap opening, most moths are not caught. Successful captures occur primarily when moths ascend into the trap after interacting with the lure, rather than retreating downwind to escape (Figure 3). Cases of escape by flying against the wind are rare. In some cases, corn earworms approach the lure, enter the trap entrance and then leave without being caught.

**Fig. 3.**
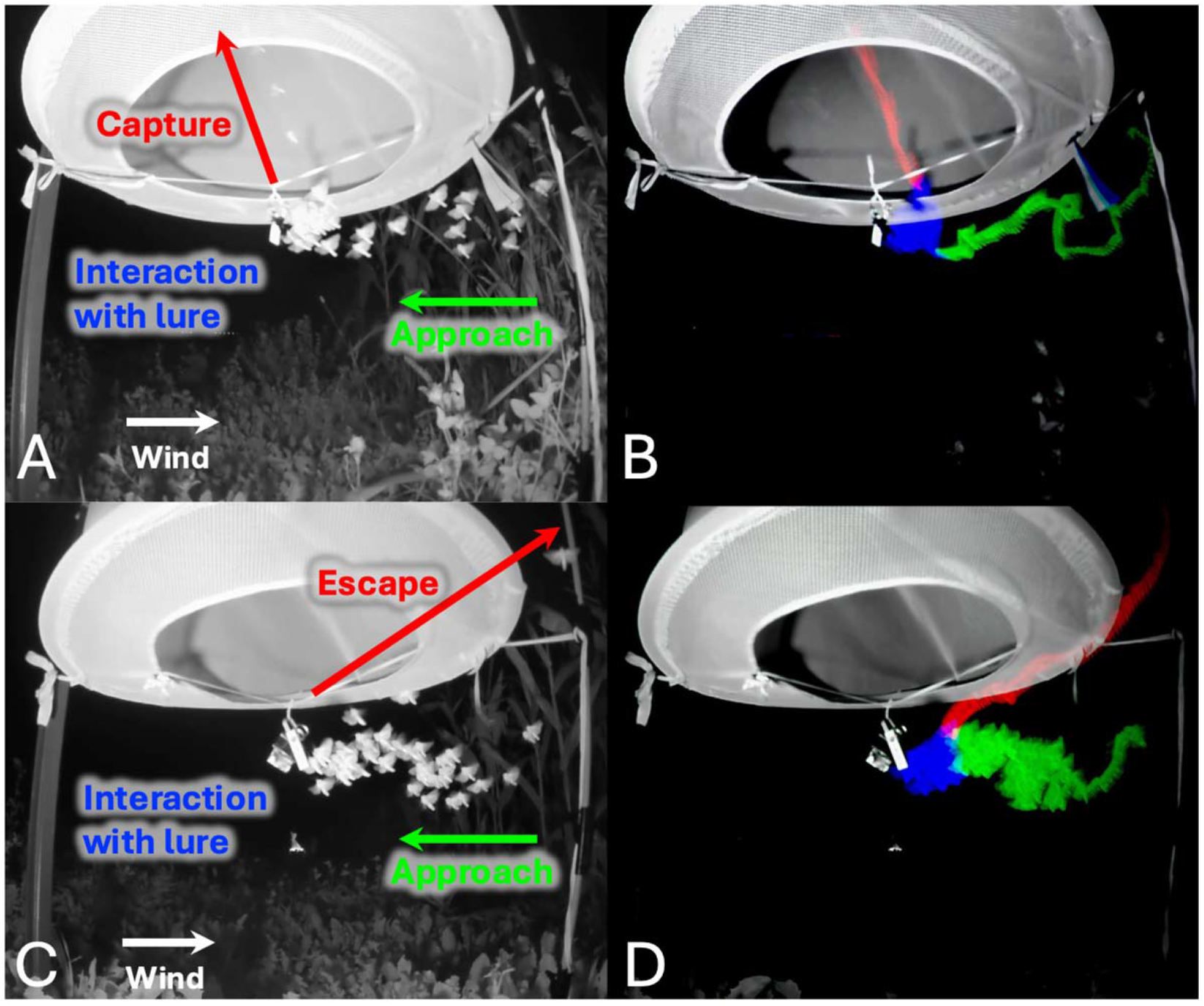
Stacked video frames showing flight behavior of corn earworm around Heliothis pheromone trap leading to capture (A) or escape (C) and the corresponding flight trajectories (B and D) illustrating approach (green), interaction with lure (blue), and escape or capture (red).

## Discussion

This study introduces a simple, affordable, and innovative approach to understanding the nocturnal flight behavior and activity of insect pests, with implications for improving trap choice, lure placement, and trap design. Using IR reflectance and IR camera, we successfully recorded the flight behavior of corn earworm, offering valuable insights into their interactions with pheromone trap and also reporting the activity time window at night. The findings have several important implications not only for corn earworm but for many different pests and trapping systems in agriculture.

For the flight behavior and trap interaction, our results reveal that a significant proportion of corn earworms approach the lure but escape without being captured. Approaches are consistently upwind, in line with the assumed diffusion of the pheromone plume in the direction of the wind. The escape behavior, often characterized by backing up downwind vertically after identifying the lure as non-female, needs further analyzed though mathematical modelization to help refine trap designs for increasing catch rate. The probability of vertical versus horizontal movement plays a critical role in determining whether moths are captured, and it appears that we have long assumed that ascending flight is the only behavior exhibited around the lure under the trap entrance.

The low catch rate of 25% on average highlights the need to optimize trap design. While the design and effectiveness of traps have been extensively studied, they have rarely been examined through the lens of insect behavior. The low catch rates observed in this study raise important questions about our ability to accurately estimate insect pest activity from trap data and how we understand trap specificity for different insect pest. For example, why are codling moths (*Cydia pomonella*) mainly caught in delta traps, whereas diamondback moths (*Plutella xylostella*) are more effectively caught in bucket traps? Similarly, the known significantly higher catch rates of corn earworms in the Hartstack trap compared to the Heliothis trap, despite their similar cone-shape design and openings, remain unanswered (Guerrero et al., 2014). A plausible explanation is that the lure in the Hartstack trap is placed closer to the entrance, possibly increasing its capture rates. Adjusting the position of the lure in the Scentry Heliothis trap to bring it closer to the entrance could improve its performance and will be tested next season. However, it is important to ensure that the pheromone plume is dispersed correctly; a significant vertical movement of the plume through the mesh trap could encourage moths to approach from the side or the top, where they could be blocked by the trap’s structure, rather than from below. Taking these nuances into account could greatly improve the effectiveness of traps and the accuracy of pest monitoring. Overall, we anticipate that a simple adjustment to the lure positioning in the Scentry Heliothis trap could enhance its performance and more competitive with the Hartstack trap which is less transportable (no possibility to fold the metal mesh trap) and harder to acquire (the original manufacturer is out of business).

This work has a broader implication for insect pest monitoring and management. The lack of comprehensive behavioral data for many nocturnal pests presents a significant challenge to IPM strategies. Therefore, we believe that developing a robust database of nocturnal insect pest behaviors is essential. Future research should investigate species-specific flight dynamics and escape mechanisms across various trap designs to better understand behavioral patterns that influence trapping success.

Efforts should also focus on optimizing trap engineering by experimenting lure placement, trap entrance shape and size, analyzing flight behavior using motion-tracking software and mathematical model, and pheromone plume direction to enhance capture rates. Interdisciplinary work between IPM specialists, chemical ecologists, behavioral entomologist, fluid physicist, and bioengineers in collaboration with the pheromone and pest management industries is essential for refining trap systems and addressing discrepancies in field monitoring. Additionally, aligning trap performance with pest activity and catch count threshold will help mitigate the underestimation of infestation levels, as frequently reported by IPM specialists, ensuring more accurate and effective pest management strategies.

In conclusion, this study demonstrates the utility of scalably affordable IR reflectance and camera in studying nocturnal insect pest behavior, providing novel insights into the design of trapping systems. By addressing our current limitations in understanding insect flight behavior around trapping systems, we can enhance pest monitoring accuracy, update catch count thresholds in pray programs, and ultimately improve precision agriculture.

## Acknowledgements

We are grateful to the Kime Farm in Geneva, NY, for generously providing access to their cornfield for this work. We also deeply appreciate the invaluable discussions on corn earworm biology and management with Michael Crossley, Galen Dively, Daniel Gilrein, Kelly Hamby, Anders Huseth, Thomas Kuhar, Brian Nault, David Owens, and everyone involved in the CEW IPM project.

## Funding

This research was funded by the U.S. Department of Agriculture, National Institute of Food and Agriculture (USDA-NIFA SCRI, grant #2023-51181-41157), the Federal Capacity Funds (Hatch Multistate Chemical Ecology NE2001 (grant #7005565), and Hatch grant #7003645).

## Author Contributions

CD (Conceptualization, Data curation, Formal analysis, Funding acquisition, Investigation, Visualization, Methodology, Writing – original draft), CR (Conceptualization, Funding acquisition, Visualization, Writing – review & editing).

## Notes

http://datadryad.org/stash/share/E78OXHMsYtNc5rTXJ5RrazKB9zQSe8vJPae8rCdCBe8

### Competing Interest Statement

The authors have declared no competing interest.

